# A visualization framework for cell division activity and orientation in pre-anthesis ovaries of *Prunus* species

**DOI:** 10.64898/2026.02.16.706041

**Authors:** Ayame Shimbo, Soichiro Nishiyama, Tatsuya Katsuno, Akane Kusumi, Hisayo Yamane, Masahiro M. Kanaoka, Ryutaro Tao

## Abstract

Fruit size and shape, which influence horticultural quality, are determined by the number and the size of the cells in the local region. In fruit trees, however, the difficulty of applying molecular genetic approaches has hindered a detailed understanding of the localization and orientation of cell division in developing fruit tissues. In this study, we established a novel framework to visualize cell division in pre-anthesis ovaries of three drupe crops, peach (*Prunus persica*), Japanese apricot (*P. mume*) and the interspecific hybrid Japanese apricot (*P. salicina* x *P. mume*), providing clear insight into the spatial distribution and orientation of dividing cells. We systematically optimized a 5-ethynyl-2′-deoxyuridine (EdU) labeling protocol for thick ovary tissues by adjusting infiltration conditions and fixation methods. In addition, electron microscopy combined with wide-view tiling visualization was applied to directly identify dividing cells, including those undergoing chromosome segregation and cell plate formation. By combining with machine learning-based detection, we efficiently and objectively identified dividing cells. Using these complementary approaches, we found that cell division activity was broadly distributed throughout pre-anthesis ovaries in all three crops, without pronounced spatial restriction. In contrast, analysis of division orientation revealed region-specific patterns: cells in the outermost exocarp divided predominantly anticlinally, whereas cells in the mesocarp divided largely periclinally, consistent with subsequent ovary (fruit) enlargement. The integrated framework presented here provides a foundation for understanding the spatial and three-dimensional regulation of fruit development and for future studies in fruit morphogenesis and horticulture.

## Introduction

The genus *Prunus*, including economically and horticulturally important fruit tree species such as peach (*Prunus persica*), Japanese apricot (*P. mume*), Japanese plum (*P. salicina*), and cherry (*P. avium*), represents a particularly significant group in fruit production. *Prunus* species characteristically produce drupes and the ovary develops into a stone fruit, in which the ovary wall differentiates into the pericarp consisting of three distinct tissue layers: the exocarp, mesocarp, and endocarp (Sterling, 1953). In fleshy-fruited *Prunus* species, the exocarp gives rise to the fruit skin, the mesocarp undergoes extensive cell proliferation and expansion to form the edible tissue, and the endocarp differentiates into a lignified stone enclosing the seed, each following distinct developmental trajectories. Most *Prunus* species have a single carpel, in which the margins grow around the central proximal-distal axis and fuse to form the suture line, corresponding to the groove on the fruit (Sterling, 1964).

Fruit size and shape are critical horticultural traits that influence consumer preference. These traits are primarily governed by cell number and cell size, highlighting the importance of elucidating the underlying cellular processes of division and expansion in fruits. Fruits are thick and three-dimensional in structure, which makes the analysis of their developmental processes challenging (Kusumi et al., 2024). However, in *Prunus*, the ovary develops into a relatively simple and rounded fruit. In addition, the relatively small genome size of *Prunus* facilitates genetic analyses related to fruit size and shape. Given the wide interspecific and intraspecific diversity in fruit size within the genus *Prunus*, as well as its horticultural importance, numerous anatomical studies have investigated differences in fruit size and shape among cultivars. For example, in peach, comparative analyses of small- and large-fruited cultivars identified cell number as a major determinant of fruit size (Ralph et al., 1991), and similar relationships between fruit size and cell number have been reported in Japanese plum and sweet cherry (Nagashima et al., 2019; Olmstead et al., 2007), suggesting that cell division plays a critical role in determining final fruit size among different cultivars. While these studies have provided valuable insights based on overall cell number or cell size, spatial information regarding where cell divisions occur within the ovary and how the orientation of cell division is regulated during the early stages of fruit formation has received less attention.

In model plants such as *Arabidopsis thaliana*, the spaciotemporal regulation of cell division has been extensively analyzed in relatively thin organs, including leaves, sepals, and petals. For example, multifaceted analyses using cell cycle reporters and staining techniques have shown that cell division in leaves does not occur uniformly across the organ, but is instead locally concentrated with high activity at the junction between the leaf blade and the petiole (Ichihashi et al., 2011). Furthermore, final organ shape has been modeled based on the position and orientation of cell divisions in primordia (Kinoshita et al., 2022). In contrast, the application of molecular genetic approaches to investigate the localization of cell division in fruit trees is technically challenging due to the long juvenile phase and the difficulty of stable transformation, particularly in *Prunus* species. Consequently, approaches relying on anatomical and microscopy-based analyses represent a feasible framework for investigating cell division in *Prunus* and other fruit tree species.

In this context, microscopy-based anatomical approaches have provided informative but still limited examples of direct visualization of cell division in fruit tissues. In the fruit of the model plant tomato (*Solanum lycopersicum*), mitotic cells have been visualized using DAPI staining, allowing detailed analyses of the orientation of cell division across different fruit regions (Renaudin et al., 2017). By contrast, in fruit trees such as *Prunus*, comprehensive analyses of cell division dynamics have not yet been achieved. Although electron microscopy allows ultrastructural observation at high resolution and mitotic figures have been observed in post-anthesis peach fruits (Masia et al., 1992), these studies did not address a detailed mapping of the spatial distribution or localization of dividing cells within the fruit tissue.

5-ethynyl-2′-deoxyuridine (EdU) labeling is a promising technique for visualizing cells undergoing division. EdU, a thymidine analog, is incorporated into nuclei of dividing cells, enabling their labeling through subsequent reaction with a fluorescent azide. It was originally developed in animal systems as a method to mark cells undergoing cell division (Salic and Mitchison, 2008), and protocols have subsequently been established for visualizing cell proliferation in plant cultured cells (Kotogány et al., 2010) and in plant leaf tissues (Nakayama et al., 2015). However, to our knowledge, the application of EdU labeling to fruit tissues has not yet been reported. Its application to fruits is expected to require methodological considerations to overcome challenges posed by the substantial thickness of fruit tissues and their strong autofluorescence, which interfere with the detection of labeled nuclei.

To address this gap, we here established an analytical framework tailored to fruit tissues by combining the optimization of EdU labeling conditions with direct observation of dividing cells using electron microscopy in pre-anthesis ovaries of *Prunus*. By systematically adjusting EdU penetration time, concentration, and fixation procedures, we achieved reliable visualization of cell division activity in thick fruit tissues. In addition, electron microscopy enabled the direct identification of cells undergoing mitosis, including those in chromosome segregation and cell plate formation. Furthermore, we introduced a machine learning–based automatic detection approach, demonstrating that an orientation-aware YOLO (You Only Look Once) model can stably, accurately, and objectively detect dividing cells even in large-scale panoramic images. We also examined the orientation of cell divisions across different regions of the ovary. Together, this study provides fundamental insights into cell division activity in *Prunus* fruits and establishes a robust experimental framework for precise analysis of cell division dynamics in drupaceous fruit tissues, thereby laying the foundation for a spatially resolved understanding of fruit development and morphology.

## Materials and methods

### Plant materials

Three drupe crops, peach ‘Akatsuki’, Japanese apricot ‘Nankou’ and the interspecific hybrid Japanese apricot ‘Tsuyuakane’ (*P. salicina* x *P. mume*), planted at the Kyoto Farmstead of the Experimental Farm of Kyoto University (Kyoto, Japan), were used for EdU labeling. Peach ‘Akatsuki’ and the interspecific hybrid Japanese apricot ‘Tsuyuakane’ were also used for electron microscopy observation. Samples were collected before flowering at BBCH scale 55 (Fig 1A; Fadón et al., 2015).

**Figure 1.**
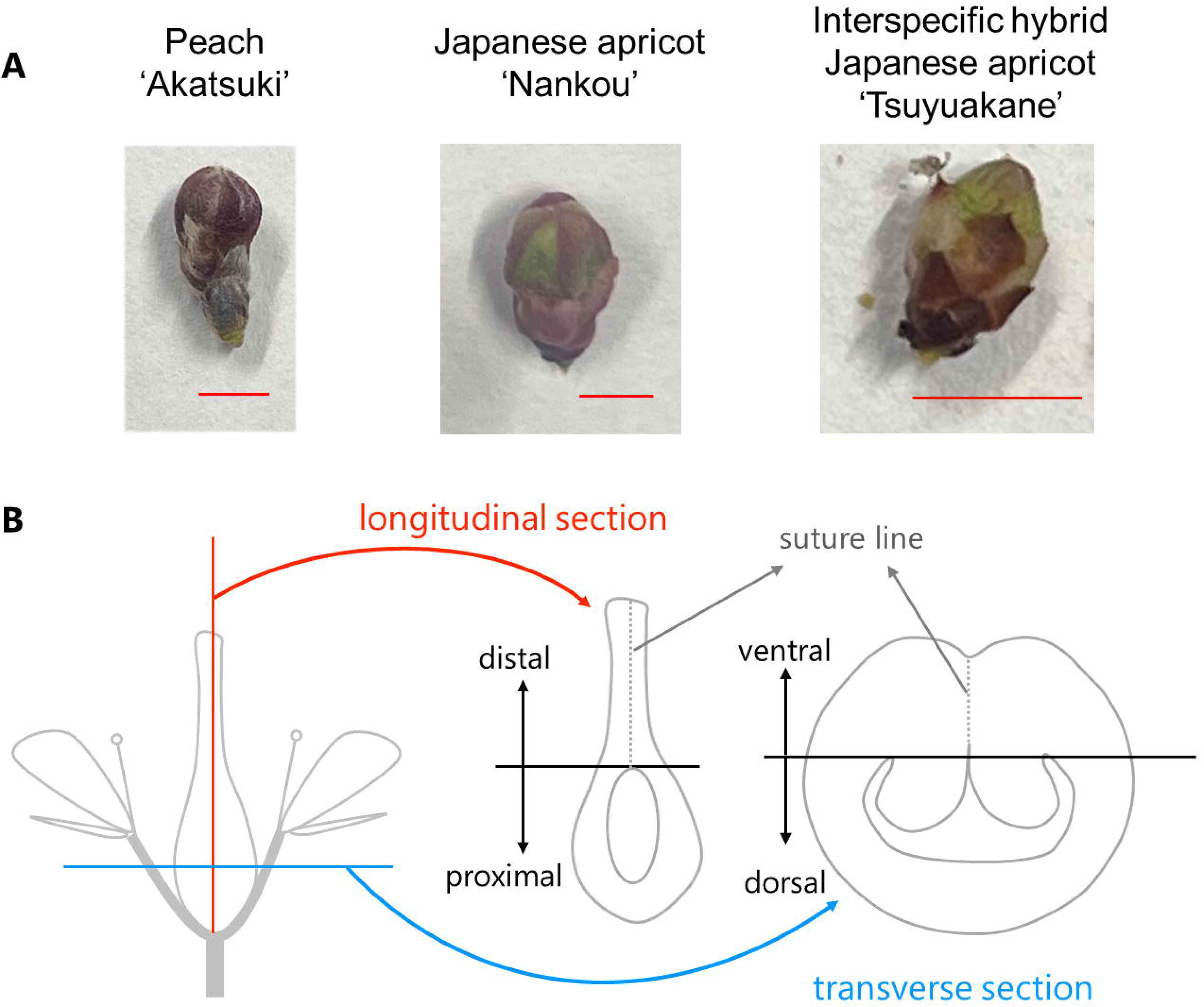
Overview of plant materials. (A) Flower buds of drupe crops at BBCH scale 55. From left to right, peach ‘Akatsuki’, Japanese apricot ‘Nankou’ and the interspecific hybrid Japanese apricot ‘Tsuyuakane’. Bar = 5 mm. (B) Schematic diagram of a drupe flower illustrating the orientation of longitudinal and transverse sections of the ovary.

### EdU labeling

EdU labeling was performed using the Click-iT™ EdU Cell Proliferation Kit for Imaging with Alexa Fluor™ 488 dye (Thermo Fisher Scientific, USA) following the manufacturer’s instructions, with modifications based on Nakayama et al. (2015). To enhance penetration of the EdU solution, ovaries were dissected to remove unnecessary outer tissues, leaving only the central region. The samples were placed in 1.5 mL tubes, and an appropriate concentration of EdU solution was added until the samples were almost completely submerged. To promote infiltration of the EdU solution, deaeration treatment and centrifugation were applied for an initial period of 30 minutes.

After EdU incorporation for an appropriate duration at room temperature, the EdU solution was removed, and 1–2 mL of 99% methanol was added for fixation. Samples were fixed on ice for 30 min and incubated at 4 °C overnight. Subsequently, the solution was replaced sequentially with sucrose solutions of 10%, 20%, and 30%, each for 2 h. Frozen sections with a thickness of 30 μm were prepared according to the Kawamoto method (Kawamoto and Shimizu, 2000). The sections were air-dried for 30 min at room temperature and then washed once with 1× PBS (phosphate-buffered saline, pH 7.4) using a pipette.

For EdU detection, a reaction cocktail was prepared immediately before use by mixing Click-iT^®^ EdU reaction buffer (227.5 μL), CuSO₄ (10 μL), Alexa Fluor^®^ 488 azide solution (0.6 μL), and Click-iT^®^ EdU buffer additive (12.5 μL) in this order. The volume of the reaction cocktail was adjusted according to the size of sections. Half of the reaction cocktail was applied onto each section using a pipette to fully cover the tissue, and the sections were incubated for 30 min at room temperature in the dark. After incubation, the solution was removed using a pipette, and the remaining half of the reaction cocktail was applied in the same manner, followed by incubation for an additional 30 min.

Following the reaction, the solution was removed, and the sections were counterstained with Hoechst33342 solution (5 μg/mL) for 30 min in the dark. Then, the sections were washed three times with 1× PBS, mounted on glass slides, and observed using a fluorescence microscope (BX53, Olympus, Japan).

### Electron microscopy

The plant materials were fixed with 2.5% (v/v) glutaraldehyde and 2% (w/v) paraformaldehyde in 100 mM phosphate buffer pH 7.2 (PB) at 4℃ overnight. After washing with 100 mM PB, the samples were post-fixed with 2% (w/v) osmium tetroxide at room temperature for 1.5 hours. Then, the samples were dehydrated in a graded series of ethanol followed by 100% propylene oxide, and embedded in epoxy resin (Luveak-812; Nacalai tesque, Japan) according to a standard procedure. The serial sections (300-400 nm thickness) were cut with a diamond knife (SYM NV3045 Ultra, SYNTEK, Japan) using an ultramicrotome (ARTOS 3D, Leica, Germany). The sections were collected on a cleaned silicon wafer strip held by a micromanipulator (MN-153, NARISHIGE, Japan). The sections were stained with 2% (w/v) aqueous uranyl acetate for 20 min and Reynolds’ lead citrate for 2 min. Backscattered electron imaging of resin-embedded sections was obtained using a scanning electron microscope (SEM) (JSM-7900F, JEOL, Japan) operated at an accelerating voltage of 6 kV, supported by Array Tomography Supporter software (version 1.3.1.0, System In Frontier Inc., Japan) that enables automated imaging. For array tomography, the images were stacked in order by Stacker NEO software (System In Frontier Inc.). Panoramic images covering the entire ovary were acquired by tile scanning at 3,000× magnification (image resolution: 2560 × 1920 pixels). The tiled images were assembled using Measurement Adviser software (System In Frontier Inc.). For ‘Tsuyuakane’, one longitudinal section and one transverse section were analyzed (Fig 1B), each obtained from a different fruit. For ‘Akatsuki’, longitudinal and transverse sections were obtained from two fruits for each orientation, with three sections analyzed per fruit.

### Machine learning

Cells undergoing chromosome segregation or cell plate formation were annotated using CVAT (version 2.43.0; https://www.cvat.ai/) with oriented bounding boxes. Annotations were performed on four panoramic images covering entire ovaries, with one longitudinal section and one transverse section from each of ‘Tsuyuakane’ and ‘Akatsuki’. All annotations were exported in the Ultralytics YOLO Oriented Bounding Box (OBB) format. In total, 280 annotated images were prepared for model training using five-fold cross-validation. In addition, 72 images confirmed by visual inspection to contain no target objects were included as negative samples in the training set of each cross-validation fold.

Object detection models were trained using YOLO11 (Jocher and Qiu, 2024). Initially, models based on conventional axis-aligned (horizontal) bounding boxes (HBB) and oriented bounding boxes (OBB) were trained on the original images with an input image size of 1920 pixels, and their detection performance was evaluated. HBB annotations were derived from the OBB annotations by calculating the minimum and maximum x–y coordinates of the annotated vertices for each object.

Based on this comparison, OBB-based models were selected, and final model training was conducted under two different training configurations. In the first configuration, patch-based training was performed with an input size of 1280 pixels to enable sliced inference using Slicing Aided Hyper Inference (SAHI; Akyon et al., 2022), which improves detection performance in large images by dividing them into overlapping patches and merging predictions from individual slices. In this configuration, defocus-like blur augmentation was applied during training to improve robustness to blur commonly encountered in microscopy images. In the second configuration, models were trained on the original images with an input image size of 1920 pixels without blur-based data augmentation. For both configurations, the models used for subsequent analyses were trained for up to 5000 epochs with early stopping enabled (patience = 500), using otherwise identical training settings. The optimizer and learning rate followed the default settings of the Ultralytics YOLO framework, with a batch size of 16.

Model performance was primarily evaluated based on recall, defined as overlap between predicted and annotated bounding boxes, reflecting the subsequent manual verification of detections by visual inspection. In addition, mAP50 and mAP50–95 were calculated as supplementary performance metrics to facilitate comparison with standard object detection benchmarks.

The trained models were applied to all original TIFF tiles comprising each panoramic ovary image. Inference was performed using a custom Python script based on the Ultralytics YOLO framework. For models trained with patch-based input, sliced inference was employed using SAHI (slice size = 1280 × 1280 pixels, overlap ratio = 0.5) to enable stable detection across the entire field of view. For models trained with an input image size of 1920 pixels, inference was performed directly on each original panoramic tile without slicing. Predictions from individual tiles were mapped back to the panoramic coordinate system using the tile coordinate metadata to reconstruct detections across the entire ovary. All predicted detections were subsequently inspected manually, and false-positive predictions were removed prior to downstream quantitative analyses. All code, annotated dataset, and the trained models have been made publicly available at https://github.com/pomology-ku/cell_div_det.

### Measurement of cell division angles

Cell division angles were measured using ImageJ (Schindelin et al., 2012) for dividing cells identified by the above machine learning–based detection. For longitudinal sections, measurements were done separately for proximal and distal regions, whereas transverse sections were divided into ventral and dorsal regions (Fig 1B). Cells were classified according to their position within the ovary into the first exocarp layer (outer epidermis, ex1), second exocarp layer (ex2), the region extending from the exocarp toward the central area (ex3), the first endocarp layer (inner epidermis, en1), the second endocarp layer (en2), and the region extending from the endocarp toward the central area (en3) (Fig 2A). In this study, ex3 and en3 were collectively defined as the mesocarp. In regions around the suture line, measurements were performed separately for three cell layers on each side of the suture line. Ovules were excluded from the analysis. Division angles in ex1–ex3 were measured relative to the exocarp surface (Fig 2B), those in en1–en3 were measured relative to the endocarp surface, and those in the suture region were measured relative to the suture line. When either the endocarp or exocarp was absent due to sectioning constraints, angles were measured relative to the remaining tissue layer. All division angles were defined within a range of 0–90 degrees.

**Figure 2.**
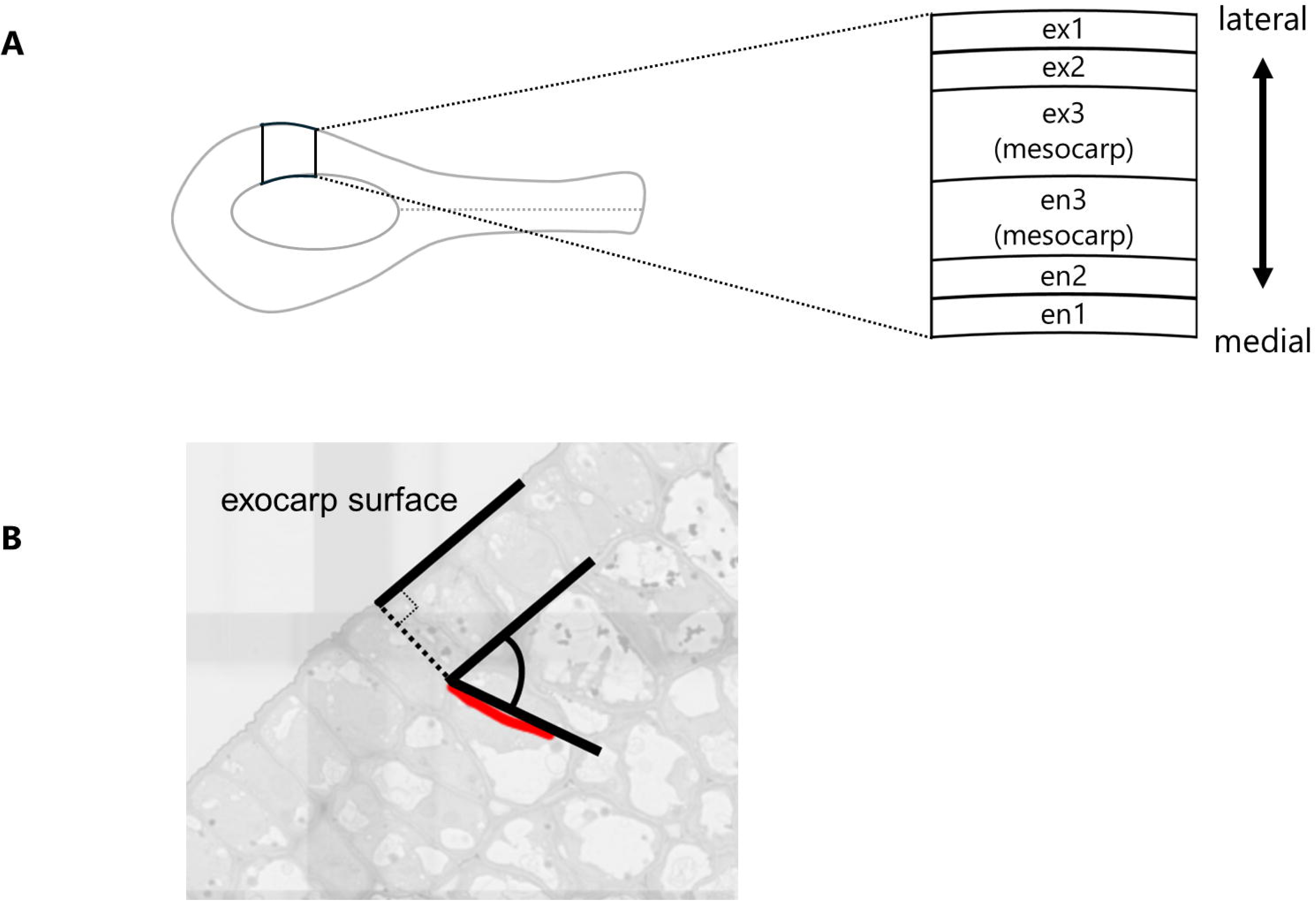
Schematic diagram of cell division angle analysis. (A) Schematic diagram illustrating the definition of cell layers within the ovary. Cells were classified into the first exocarp layer (ex1), second exocarp layer (ex2), the region extending from the exocarp toward the central area (ex3), the first endocarp layer (en1), the second endocarp layer (en2), and the region extending from the endocarp toward the central area (en3). (B) Schematic illustration of angle measurement. Angles were measured relative to the pericarp surface. Division angles were defined within a range of 0–90 degrees.

## Results

### Optimization of EdU labeling for ovary tissues

Protocols for EdU labeling have been established for leaf tissues (Nakayama et al., 2015); however, their direct application to fruit tissues has not been fully evaluated. Therefore, in this study, we optimized EdU labeling conditions for pre-anthesis ovaries by examining EdU concentration, incubation time, and fixation methods. In tomato leaves, EdU labeling at 10 µM for 2–3 h has been shown to provide reliable detection of proliferating cells (Nakayama et al., 2015). In drupe developing ovaries, we found through empirical testing that higher EdU concentrations (25–100 µM) combined with longer incubation periods (8–10 h) were required to obtain robust and reproducible fluorescence signals.

Fixation conditions were also critical for successful detection. In leaf tissues, fixation with FAA for 2–3 h has been used (Nakayama et al., 2015); when applied to fruit tissues, this procedure resulted in strong autofluorescence that interfered with the EdU signal observation. By contrast, overnight fixation with methanol effectively reduced autofluorescence and markedly improved the clarity of EdU fluorescence signals in drupe developing ovaries.

Using the optimized protocol, clear EdU signals were successfully detected in ovaries of three drupe species, ‘Akatsuki’, ‘Nankou’, and ‘Tsuyuakane’ (Fig 3A–C). Counterstaining with Hoechst 33342 revealed that all nuclei were labeled by Hoechst 33342, whereas EdU signals were detected only in a subset of nuclei, suggesting that EdU incorporation occurred specifically in nuclei undergoing DNA replication (Fig 3D).

**Figure 3.**
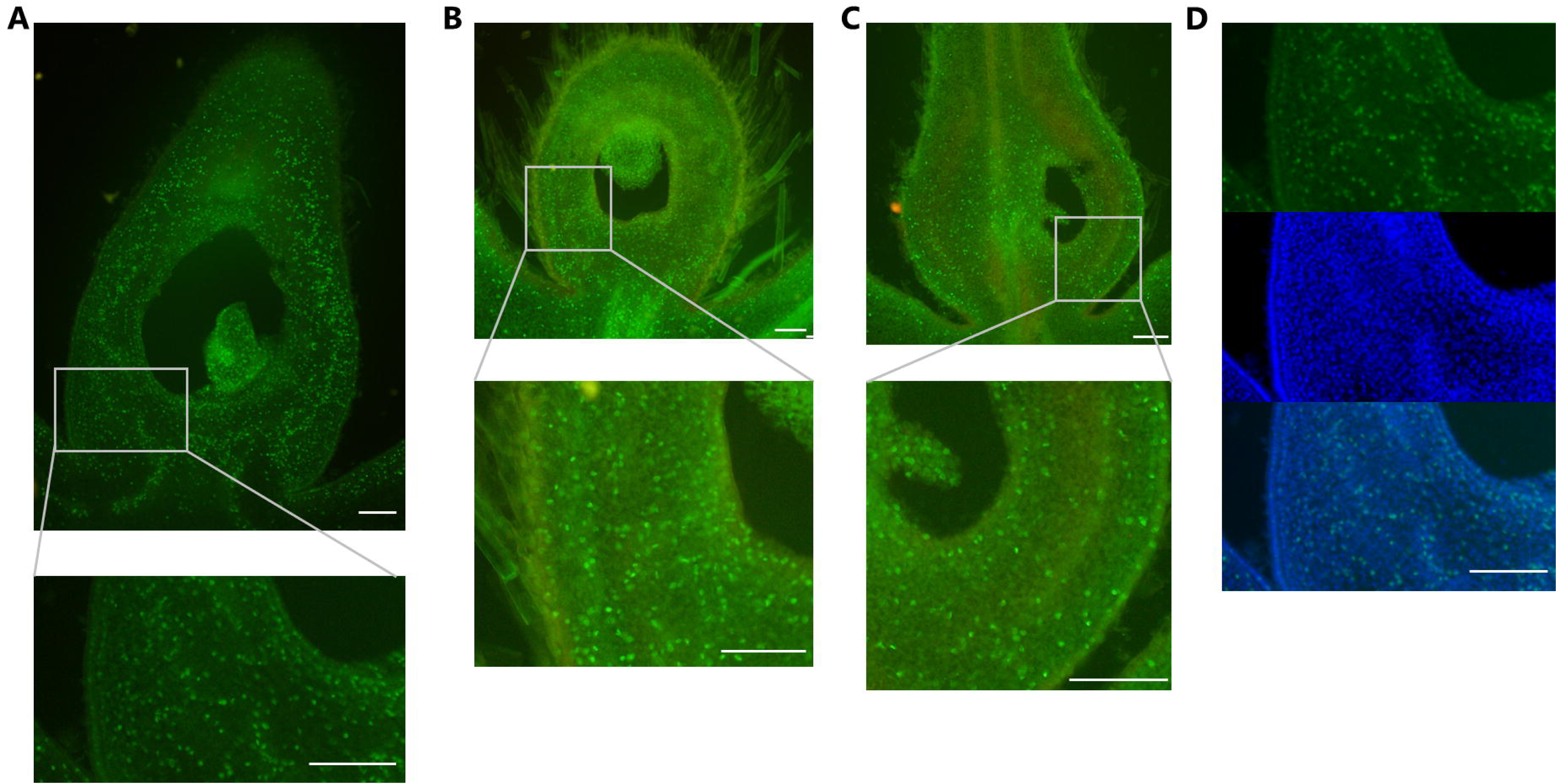
Visualization of EdU incorporation in pre-anthesis ovaries. (A) Peach ‘Akatsuki’. EdU concentration was 50 µM, and the infiltration period was 8 h. (B) Japanese apricot ‘Nankou’. EdU concentration was 25 µM, and the infiltration period was 8 h. (C) The interspecific hybrid Japanese apricot ‘Tsuyuakane’. EdU concentration was 100 µM, and the infiltration period was 10 h. Bottom panels in (A) to (C) are the magnified images of square regions in the top panels, respectively. (D) Representative nuclear signals showing EdU labeling (top), Hoechst 33342 staining (middle), and merged (bottom) images. Bars = 100 µm.

This approach enabled relatively rapid visualization of the spatial distribution of dividing cells in pre-anthesis ovaries. In drupe ovaries before flowering, EdU-positive signals were broadly distributed and appeared scattered throughout the tissue, without clear spatial localization. A similar scattered pattern of cell division was observed across all three drupe crops examined (Fig 3).

### Observation of dividing cells using electron microscopy

To complement EdU-based detection, we employed electron microscopy to directly observe dividing cells. Ultrathin sections by SEM enabled clear visualization of cells at specific mitotic stages, including anaphase cells showing chromosome segregation (Fig 4A) and telophase cells undergoing cell plate formation (Fig 4B). In addition to stage-specific identification, electron microscopy allows direct measurement of cell division orientation based on ultrastructural features.

**Figure 4.**
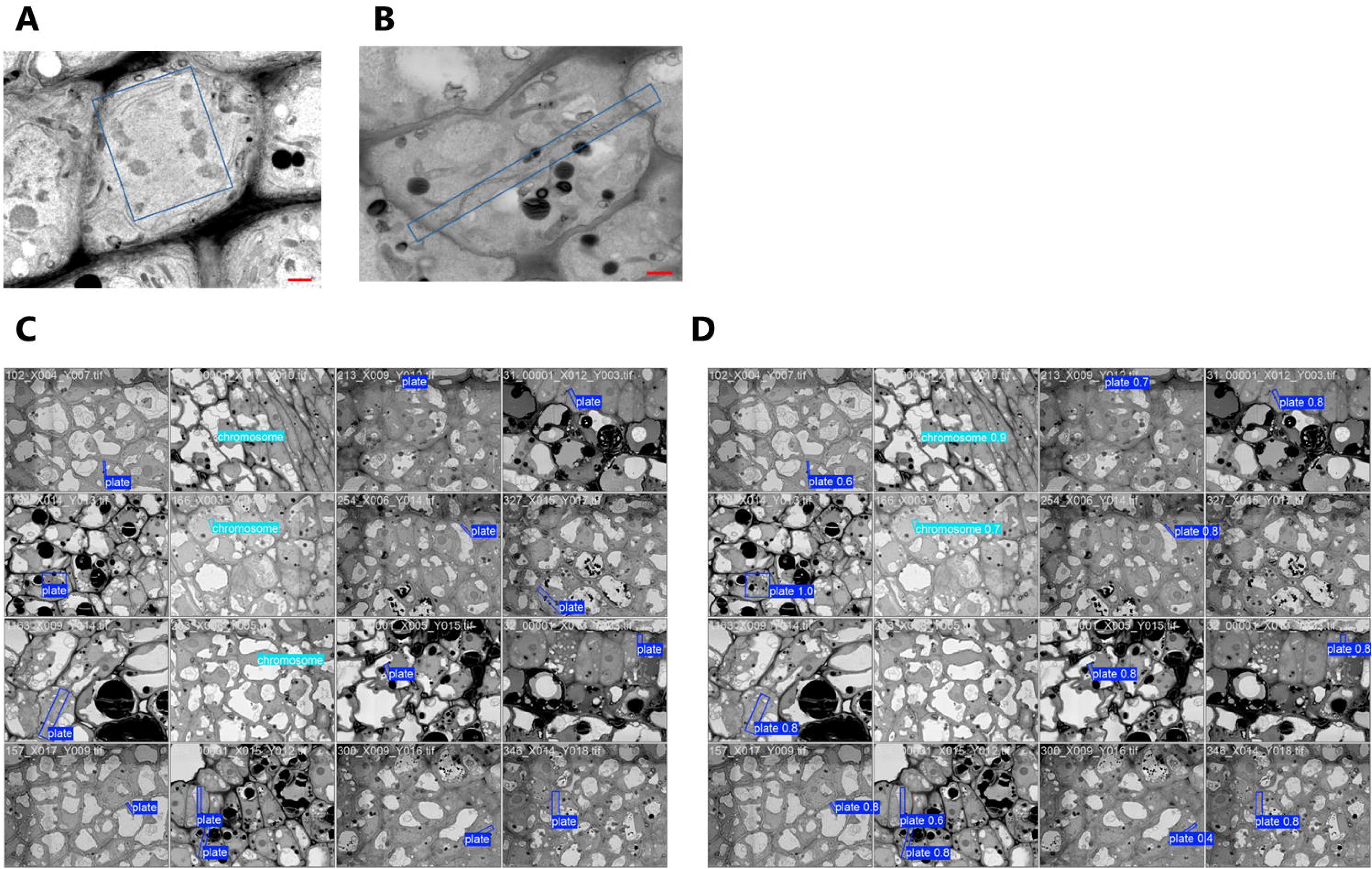
Electron microscopic visualization of dividing cells. (A) Representative cell undergoing chromosome segregation, with blue squares indicating annotated regions. Bar = 1 µm. (B) Representative cell undergoing cell plate formation, with blue squares indicating annotated regions. Bar = 1 µm. (C) Example of manual annotations applied to the electron micrograph. (D) Corresponding machine learning-based detection results obtained using the YOLO11l-OBB model.

Serial-section analysis through array tomography (Micheva and Smith, 2007) revealed that cells undergoing cell plate formation were more frequently and consistently detected than cells undergoing chromosome segregation. In cell plate–forming cells, 26 out of 28 serial sections per cell displayed clear cell plate structures (Supplementary Fig S1), whereas in cells undergoing chromosome segregation, only 8 out of 22 sections per cell exhibited recognizable mitotic features (Supplementary Fig S2). The extended persistence and distinct ultrastructural features of the cell plate formation stage likely account for its higher detectability.

### Machine learning-based detection of dividing cells

Manual identification of dividing cells in large ovary sections is highly labor-intensive and may be prone to oversight due to the large number of cells present. To facilitate a more efficient and objective analysis, we applied a machine learning–based approach for automated detection of dividing cells. HBB-based models showed limited detection performance, with mAP50–95 values below 0.07 and low recall (Table 1). By contrast, detection performance was markedly improved when OBB were used. Across all model sizes, OBB-based models achieved substantially higher mAP and recall values, with mAP50 exceeding 0.76 and mAP50–95 values in the range of approximately 0.39– 0.46 (Table 2). PR curves demonstrated consistently improved recall across confidence thresholds for OBB-based models (Supplementary Fig S3), reaching approximately 0.75 at the largest model evaluated (Table 2). These results indicate that orientation-aware detection substantially improves both the accuracy and sensitivity of detecting dividing cells in fruit tissues.

**Table 1.** Performance of horizontal bounding box detection models. Values are reported as mean ± SD over five-fold cross-validation runs. All models were trained for 100 epochs. mAP50 and mAP50–95 correspond to the best-performing epoch of each run, whereas recall is reported from the final (last) epoch. IoU was computed using the default Ultralytics YOLO implementation for each task, and mAP50–95 was averaged over IoU thresholds ranging from 0.50 to 0.95.

| model | task | image size | mAP50 | mAP50-95 | last Recall |
| --- | --- | --- | --- | --- | --- |
| yolo11n | det | 1920 | $0.1550 \pm 0.0268$ | $0.0630 \pm 0.0135$ | $0.2442 \pm 0.0775$ |
| yolo11s | det | 1920 | $0.0574 \pm 0.0394$ | $0.0215 \pm 0.0202$ | $0.0957 \pm 0.0407$ |
| yolo11m | det | 1920 | $0.0375 \pm 0.0266$ | $0.0129 \pm 0.0099$ | $0.2849 \pm 0.2173$ |

Values are reported as mean ± SD over five-fold cross-validation runs. All models were trained for 100 epochs. mAP50 and mAP50-95 correspond to the best-performing epoch of each run, whereas recall is reported from the final (last) epoch. IoU was computed using the default Ultralytics YOLO implementation for each task, and mAP50-95 was averaged over IoU thresholds ranging from 0.50 to 0.95.

**Table 2.**
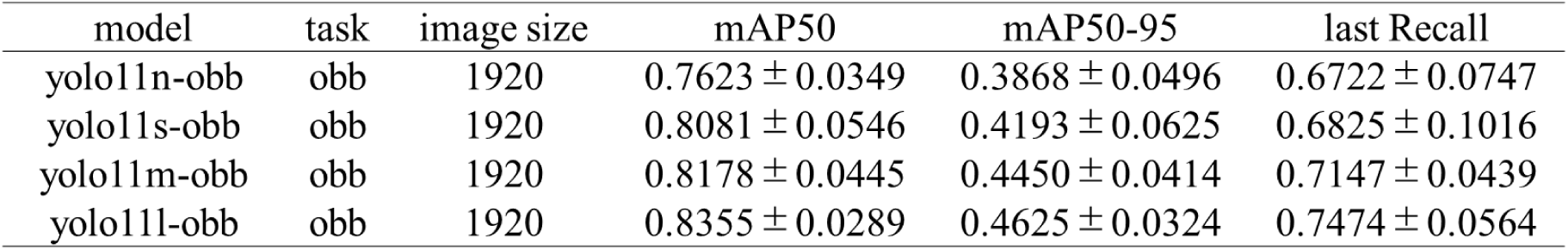
Performance of oriented bounding box detection models. Values are reported as mean ± SD over five-fold cross-validation runs. All models were trained for 100 epochs. mAP50 and mAP50–95 correspond to the best-performing epoch of each run, whereas recall is reported from the final (last) epoch. IoU was computed using the default Ultralytics YOLO implementation for each task, and mAP50–95 was averaged over IoU thresholds ranging from 0.50 to 0.95.

Based on these comparisons, two OBB-based models with partially complementary detection characteristics were applied to large panoramic images (Table 3). Qualitative inspection indicated that the two models detected overlapping but non-identical subsets of dividing cells, and combining their detections (union of detections) increased recall to 0.95. Representative detection results are shown in Fig 4C and D.

**Table 3.**
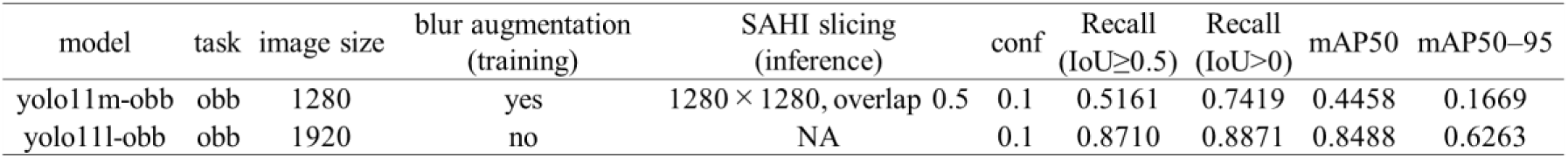
Validation performance of OBB models with different training and inference strategies. Metrics were evaluated on the validation set at the best-performing epoch. The two OBB models were trained with different augmentation strategies and inference settings to capture complementary detection characteristics. mAP50 and mAP50–95 were computed using rotated bounding box IoU (polygon IoU) over IoU thresholds from 0.50 to 0.95. Recall was evaluated at conf = 0.1 using two criteria: IoU ≥ 0.5 and any spatial overlap (IoU > 0), the latter reflecting subsequent manual curation of detections.

Using the two complementary models, dividing cells were detected across the entire ovary of ‘Akatsuki’, and their spatial distribution after manual removal of false-positive detections is shown in Fig 5, with the longitudinal and transverse sections presented in panels A and B, respectively. Cells undergoing cell plate formation were more frequently detected than cells exhibiting chromosome segregation. Consistent with the EdU labeling results, dividing cells were broadly distributed throughout the ovary. This distribution pattern was observed consistently in both longitudinal and transverse sections, indicating that cell division occurs across the entire ovary rather than being restricted to specific regions.

**Figure 5.**
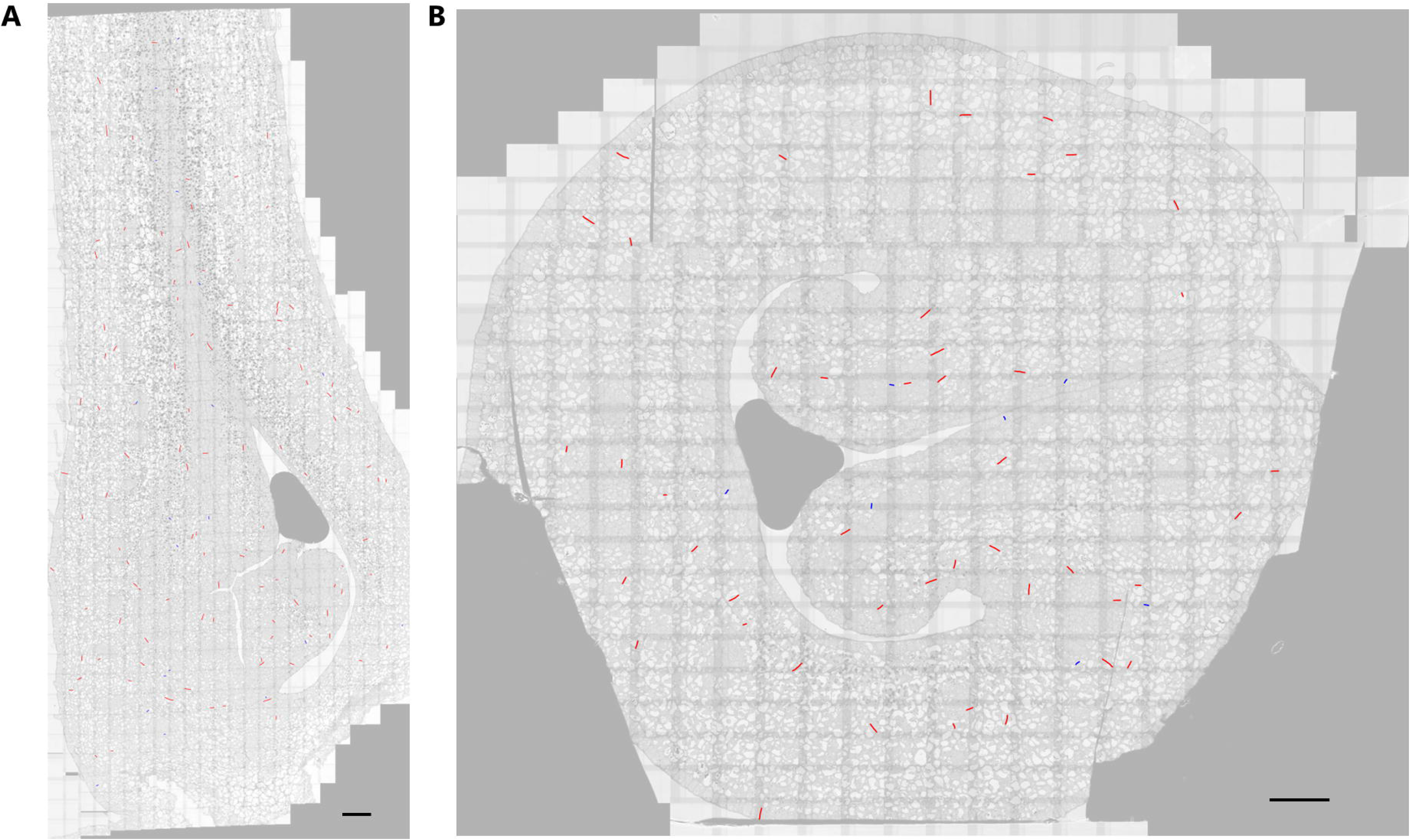
Spatial distribution and orientation of cell division in the ovary. Dividing cells detected by machine learning models and curated by manual removal of false positives are shown. Red lines indicate cells undergoing cell plate formation, and blue lines indicate cells undergoing chromosome segregation. Lines indicate the orientation of cell division. Bar = 50 µm. (A) Longitudinal section of ‘Akatsuki’. (B) Transverse section of ‘Akatsuki’.

### Analysis of cell division angles

In sections with dividing cells identified using these detection models, analysis of division angles revealed distinct, layer-dependent patterns throughout the ovary in both longitudinal (Fig 6A) and transverse sections (Fig 6B). Overall, division orientation depended more strongly on tissue layer than on positional differences along the proximal–distal or ventral–dorsal axes.

**Figure 6.**
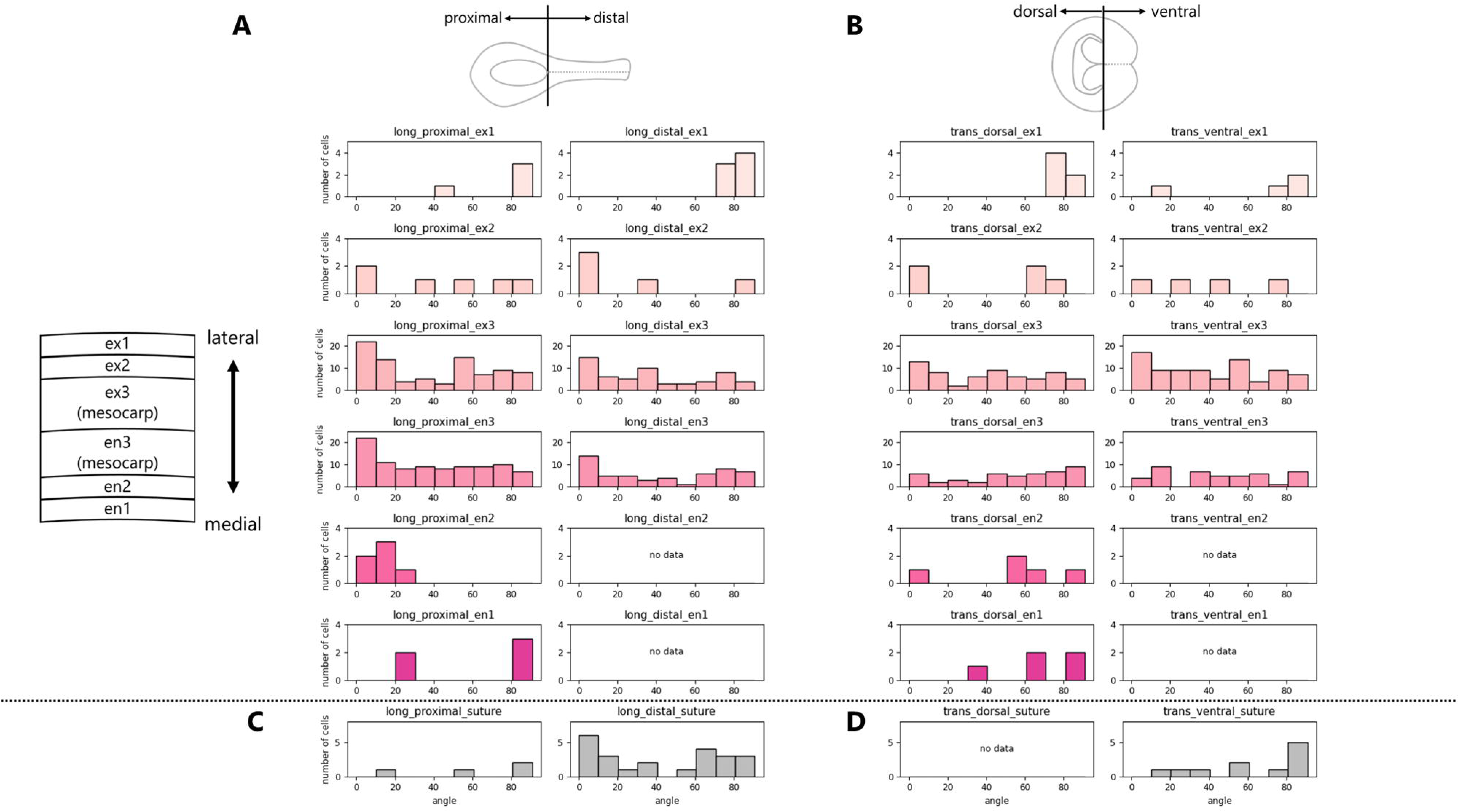
Distribution of cell division orientations in ovary tissues. For each graph, the x-axis represents division angles binned at 10-degree intervals, and the y-axis represents the number of dividing cells. (A) Distribution of cell division angles in longitudinal sections for each region. Results are shown separately for proximal (left) and distal (right) regions. (B) Distribution of cell division angles in transverse sections for each region. Results are shown for dorsal (left) and ventral (right) regions. In panel (A) and (B), results are presented from top to bottom, ex1, ex2, ex3, en3, en2, and en1. (C) Distribution of cell division angles in longitudinal sections for suture region. (D) Distribution of cell division angles in transverse sections for suture region.

In ex1 (first layer of exocarp), cell divisions predominantly exhibited anticlinal orientations (perpendicular to the tissue surface) in both the proximal and distal regions of longitudinal sections, as well as in the ventral and dorsal regions of transverse sections. Notably, this orientation bias was specific to the ex1 layer and was not observed in the adjacent ex2 layer or the innermost en1 layer.

By contrast, divisions in the more internal layers, ex3 and en3, tended to be biased toward periclinal orientations (parallel to the tissue surface), with the exception of en3 in transverse sections. In the en2 layer, no clear orientation preference was observed in transverse sections, whereas longitudinal sections showed a predominance of anticlinal divisions. In regions surrounding the suture line, division orientations were section-dependent; in longitudinal sections, no clear preferential orientation was observed (Fig 6C), whereas in transverse sections, divisions were more frequently anticlinal relative to the suture line (Fig 6D).

Spatial comparisons along the proximal–distal regions in longitudinal sections revealed no substantial differences in division orientation; similarly, in transverse sections, division angle distributions were broadly similar between the ventral and dorsal regions.

## Discussion

In this study, we developed an experimental framework that enables high-resolution visualization of cell division in the developing ovaries of *Prunus* by optimizing EdU labeling and combining it with complementary electron microscopy observations. The results obtained from EdU labeling and electron microscopy were consistent, revealing that dividing cells are broadly and randomly distributed throughout the pre-anthesis ovary. Building on protocols established for thin leaf tissues (Nakayama et al., 2015), optimization of fixation methods, EdU concentration, and infiltration time allowed clear detection of labeled-nuclei in developing ovaries.

EdU labeling provides a simple, rapid, and highly practical approach for visualizing the spatial distribution of dividing cells in pre-anthesis ovaries. This method is also applicable to relatively large samples to simply investigate the spatiotemporal localization of cell division. In fruit trees, the application of molecular genetic approaches, such as cyclin–GUS reporters widely used in model plants, is often constrained by difficulties in genetic transformation, and therefore EdU labeling serves as a useful and effective alternative for detecting proliferative activity without genetic modification. However, accurate application requires careful calibration of several parameters, including EdU concentration and incubation duration, and reliable detection depends on sufficient tissue penetration. Additionally, EdU may have potential harmful effects on cells and DNA (Zhao et al., 2013). Thus, the use of EdU labeling in conjunction with complementary techniques, such as electron microscopy, would provide a more robust framework for assessing cell division activity.

Electron microscopy further enabled direct observation of cells undergoing mitosis, including stages of chromosome segregation and cell plate formation, providing unequivocal identification of cells undergoing mitosis based on ultrastructural characteristics. On the other hand, electron microscopy demands labor-intensive preparation of ultrathin sections, and the lack of intrinsic fluorescence necessitates laborious identification of dividing cells. In this study, the latter limitation was mitigated through the application of machine learning-based detection, which enabled efficient and objective identification of dividing cells even in whole ovary sections that contain numerous cells. Similar to recent applications of machine learning to microscopic images for detecting specific cellular components, such as chloroplasts in plant tissues (Su et al., 2025), the integration of microscopy and deep learning thus provides an effective framework for spatial analysis of cell division in complex fruit tissues.

Understanding the orientation of cell divisions is essential for elucidating the mechanisms underlying fruit development. However, many previous studies on fruit size have largely focused on measurements of cell number and cell size (Olmstead et al., 2007; Ralph et al., 1991), with comparatively less attention given to the spatial and orientational aspects of cell division. This is largely due to technical challenges associated with applying molecular genetic approaches in fruit trees, such as KNOLLE–GFP reporters, in which the syntaxin KNOLLE plays an essential role in cell plate formation (Reichardt et al., 2007). In *Prunus*, only a limited number of microscopy-based studies have provided direct observations of cell division, including the valuable work of Masia et al. (1992) in peach, which used transmission electron microscopy to examine cellular ultrastructure. In this study, we expand upon these previous observations by visualizing cell division using two complementary approaches, enabling analyses that resolve both spatial distribution and division orientation. In the first exocarp layer (ex1), anticlinal divisions were frequently observed in both longitudinal and transverse sections (Fig 6A, B). This pattern is consistent with a potential contribution to isotropic surface expansion during fruit growth. A predominance of anticlinal divisions in the epidermis has been reported in tomato (Renaudin et al., 2017) and grape (*Vitis vinifera*; Considine and Knox, 1981), suggesting that abundant anticlinal divisions in ex1 may represent a conserved feature of fruit development across species. Notably, in *Prunus* ovaries, this anticlinal bias was restricted to the first exocarp layer and was not observed in the immediately underlying layers (ex2; Fig 6A, B). By contrast, in ex3 and en3, corresponding to the mesocarp, cell divisions often proceed in periclinal orientations, which may be associated with tissue thickening of the fruit flesh during development. Interestingly, in regions surrounding the suture line, anticlinal divisions were more frequently observed in transverse sections, suggesting the presence of a structural axis centered on the suture. Because the suture line is formed at the margins of carpel fusion (Sterling, 1964) and is derived from outer pericarp tissues, the persistence of anticlinal divisions in this region may reflect its developmental origin.

The results of EdU labeling and complementary SEM observations revealed widespread distribution of cell divisions in the pre-anthesis ovaries across diverse *Prunus* species. In our previous three-dimensional analyses of drupe fruit development, which focused on stages from approximately 1-2 months after anthesis through fruit maturation, growth activity was higher in the proximal region than in the distal region (Shimbo et al., 2026). A similar proximal-dominant growth pattern has also been reported in persimmon (*Diospyros kaki*) (Kusumi et al., 2024; 2025), suggesting that spatial heterogeneity in fruit growth may be a common feature among fleshy fruits. Based on these observations, we initially hypothesized that cell division might be spatially biased to the proximal region at the pre-anthesis stage. However, in the present study, cell division was observed throughout the ovary, without any obvious spatial localization at this early developmental stage. This difference likely reflects stage-specific regulation of growth processes. At the pre-anthesis stage, cell division may occur relatively uniformly across the ovary to establish a foundational cellular framework, whereas spatial heterogeneity in fruit growth observed after anthesis may arise predominantly from differential cell expansion and/or changes in the orientation or rate of cell division during later developmental stages. In drupe fruits, the ovary develops as the fruit, whereas in pome fruits of the same Rosaceae family, such as apples (*Malus* x *domestica*) and pears (*Pyrus* spp.), the hypanthium develops as the fruit, indicating differences in cell division distributions between drupes and pomes. Moreover, even within pomes, the European pear (*Pyrus communis*), which has a unique fruit shape, may exhibit species-specific cell division patterns. Furthermore, Masia et al. (1992) reported that in peach, the orientation of cell divisions in the epicarp shifts during development, initially anticlinal and later becoming multidirectional. How division orientation changes during development, and how such transitions are controlled, remain unresolved questions for future investigation. The two complementary approaches established in this study provide a foundation for addressing these questions and are anticipated to be broadly applicable across diverse species and developmental stages in future investigations.

## Conclusion

In this study, we established an anatomical framework to visualize cell division in fruit tissues by integrating EdU labeling with complementary electron microscopy. EdU labeling offers a comparatively labor-efficient approach to assess the localization of dividing cells, and the results suggested that in *Prunus* ovaries before anthesis, cell division occurs broadly throughout the ovary without apparent spatial localization. To further examine this pattern, we analyzed electron micrographs using machine learning-based detection, which enabled efficient and objective identification of dividing cells in large ovary sections. The spatial pattern observed through EdU labeling was independently supported by electron microscopy-based analysis. Ultrastructural observation also enabled clear identification of cells undergoing division, including chromosome segregation and cell plate formation, and allowed direct analysis of the orientation of cell division. We found that cells in the outermost exocarp predominantly divide anticlinally relative to the surface, whereas cells in the mesocarp largely show periclinal divisions. Together, these approaches establish a new experimental system capable of visualizing the spatial patterns of cell division in fruit floral tissues, providing a versatile foundation for future studies on fruit development and morphogenesis.

## Acknowledgment

This research was supported by the Japan Society for the Promotion of Science KAKENHI (grant no. 21KK0269 to SN, no. 24KJ1497 to AK and no. 24H00510 to SN, MMK and RT), the JST BOOST Program (grant no. JPMJBY24F7 to SN) and a research grant from the Hirose Foundation to SN.

## Author Contributions

SN conceived and designed the study. AS collected the plant materials and acquired the data. TK optimized the electron microscopy protocols, prepared specimens, and assisted with image acquisition. AK and MMK contributed to experimental design, EdU visualization, and data interpretation. AS and SN developed the methodology, analyzed the data, and drafted the manuscript. SN, HY and RT prepared plant materials and experimental facilities. All authors read and approved the manuscript.

