## Supplementary Materials for "A visualization framework for cell division activity and orientation in pre-anthesis ovaries of *Prunus* species"

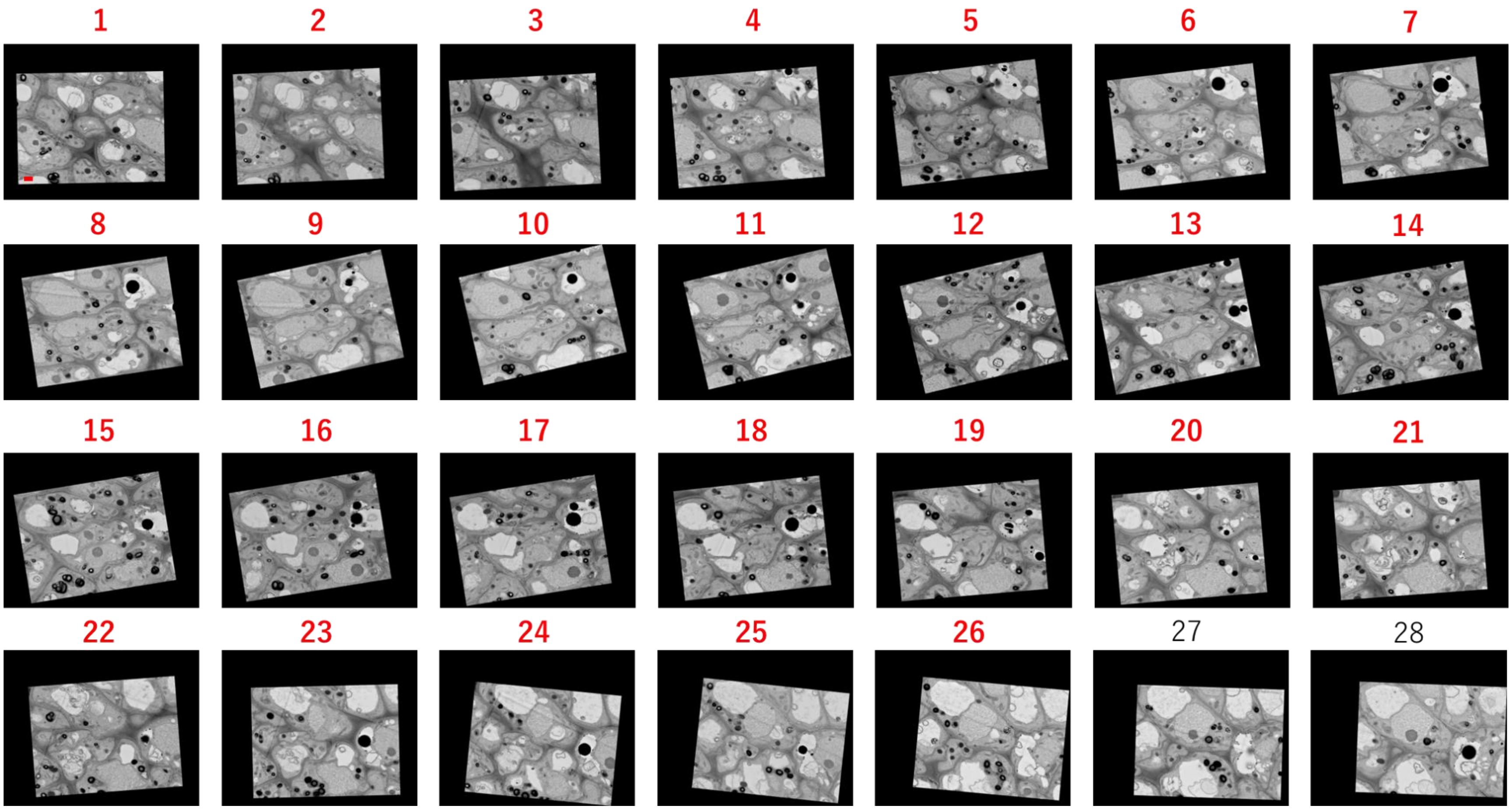


**Supplemental figure S1. Array tomography of a representative cell undergoing cell plate formation.**
Serial images along the z-axis show the same cell across different z-axis positions. Of the 28 serial sections, cell plates were observed in sections 1–26, which are highlighted in red. Bar = 1 μm.

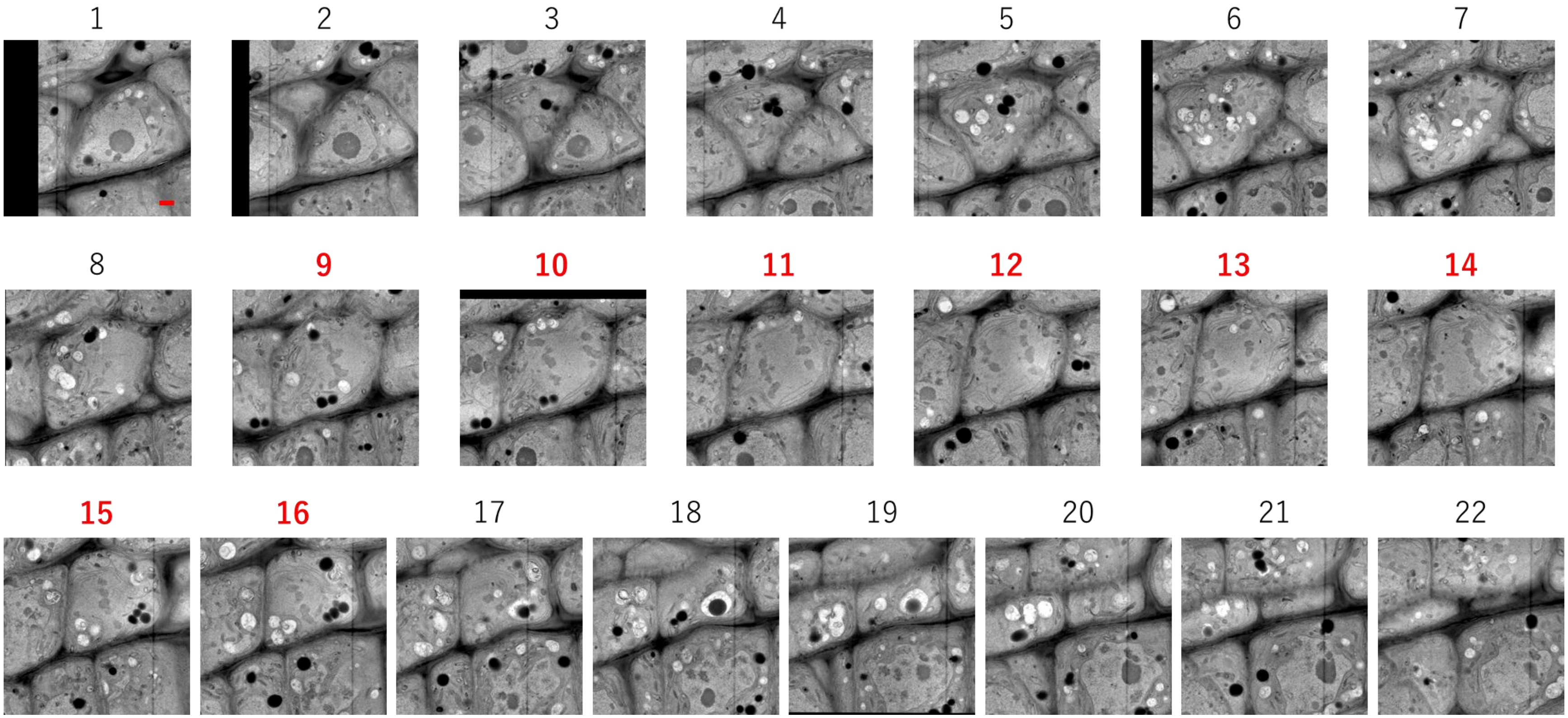


**Supplemental figure S2. Array tomography of a representative cell undergoing chromosome segregation.**
Serial images along the z-axis show the same cell across different z-axis positions. Of the 22 serial sections, chromosome segregation was observed in sections 9-16, which are highlighted in red. Bar = 1 µm.

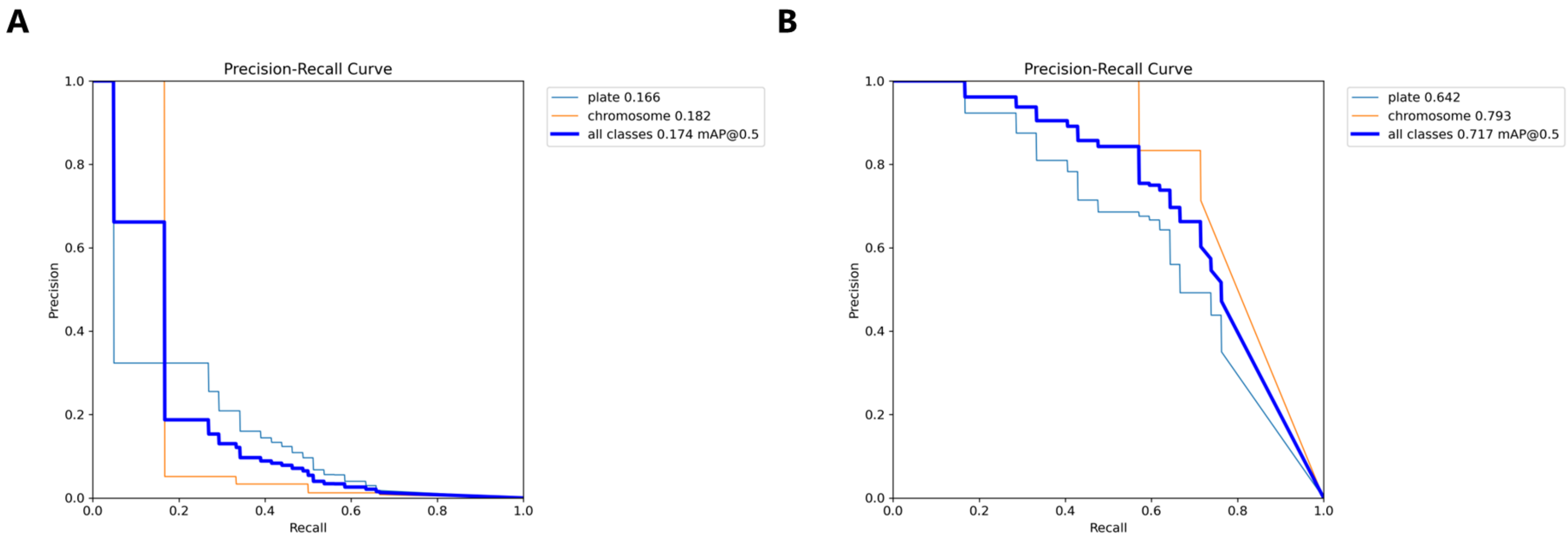


**Supplementary Fig S3. Precision–recall curves for the trained YOLO11n detection models.** Precision–recall (PR) curves obtained from a single validation fold of the YOLO11n model for (A) the horizontal bounding box (HBB) task and (B) the oriented bounding box (OBB) task.
